# Chemical mutagenesis of *Listeria monocytogenes* for increased tolerance to benzalkonium chloride shows independent genetic underpinnings and off-target antibiotic resistance

**DOI:** 10.1101/2023.12.11.571107

**Authors:** Tyler D. Bechtel, Julia Hershelman, Mrinalini Ghoshal, Lynne McLandsborough, John G. Gibbons

## Abstract

*Listeria monocytogenes*, a potentially fatal foodborne pathogen commonly found in food processing facilities, creates a significant economic burden that totals more than $2 billion annually in the United States due to outbreaks. Quaternary ammonium compounds (QACs), including benzalkonium chloride (BAC), are among the most widely used sanitizers to inhibit the growth and spread of *L. monocytogenes* from food processing facilities. However, resistance to QACs has been increasing in *L. monocytogenes* and different genetic mechanisms conferring resistance have been discovered. Here, we used ethyl methanesulfonate (EMS) to chemically mutagenize the BAC-susceptible strain, *L. monocytogenes* FSL-N1-304. We isolated two mutants with increased tolerance to BAC compared to the parental strain. Next, we assessed the off-target effect of increased tolerance to BAC by measuring the minimum inhibitory concentrations (MICs) of a diverse set of antibiotics, revealing that *mut-1* and *mut-2* displayed significantly increased resistance to fluoroquinolone antibiotics compared to the parental strain. We then sequenced the genomes of the parental strain and both mutants to identify mutations that may be involved in the increased resistance to BAC. We identified 3 and 29 mutations in *mut-1* and *mut-2*, respectively. *mut-1* contained nonsynonymous mutations in *dagK* (a diacylglycerol kinase), lmo2768 (a permease-encoding gene), and *lmo0186* (resuscitation promoting factor). *mut-2* contained a nonsense mutation in the nucleotide excision repair enzyme UvrABC system protein B encoding gene, *uvrB*, which likely accounts for the higher number of mutations observed. Transcriptome analysis in the presence of BAC revealed that genes related to the phosphotransferase system and internalins were upregulated in both mutants, suggesting their significance in the BAC stress response. These two mutants provide insights into alternative mechanisms for increased BAC tolerance and could further our understanding of how *L. monocytogenes* persists in the food processing environment.

## Introduction

*Listeria monocytogenes* is a gram-positive, foodborne bacterial pathogen that causes the serious infection listeriosis. Listeriosis is a relatively rare disease, but it can lead to severe and potentially fatal symptoms for elderly and immunocompromised individuals including infections resulting in meningitis and septicemia [1,2]. Additionally, pregnant women are more susceptible to listeriosis, which is particularly problematic as *L. monocytogenes* possesses the unique ability to permeate the placenta and infect the fetus, which can lead to fetal meningitis and miscarriages [3]. A study analyzing data from 2000-2008 revealed that *L. monocytogenes* had the highest hospitalization rate (94%) and third highest fatality rate (15.9%) among 31 major foodborne pathogens [4].

*L. monocytogenes* is highly abundant in the natural environment and capable of surviving adverse conditions, resulting in frequent contamination and persistence of *L. monocytogenes* strains infood processing facilities [5]. Given the health concerns and economic impacts of *L. monocytogenes* contamination in the food industry, the FDA has established guidelines to prevent contamination and outbreaks in foods [6]. These guidelines include personal protective equipment for workers, structured operation plans of the plant, specific equipment, testing protocols, and sanitization. Quaternary ammonium compounds (QACs) are a class of antimicrobial compounds commonly used for surface sanitization in food processing facilities [7]. The bactericidal mode of action for QACs is similar to that of a detergent. The negatively charged bacterial cellular membrane interacts with the positively charged head of the QAC molecule, allowing the nonpolar QAC side chains to disrupt the intramembrane region. This interaction leads to the formation of micelles, cytosolic leakage, and eventual cellular lysis [8].

Benzalkonium chloride (BAC) is a type of QAC that is a common component of commercial disinfectants that are used to sanitize solid surfaces in food processing facilities [7,9]. Over the last two decades, several studies have reported increased BAC tolerance in *L. monocytogenes* strains isolated from food environments [10–15]. An increase in QAC tolerance has been attributed to improper application, dilution, and biodegradation of QACs resulting in sub-lethal concentration gradients [16,17]. Several mechanisms for BAC resistance have been discovered. Most commonly, BAC resistance, or increased tolerance, involves alterations to efflux pumps [11,13,18–20]. Adaptation to BAC can also result in cross-tolerance to fluoroquinolone antibiotics, suggesting shared resistance mechanisms [21–24]. Thus, the improper usage of BAC, and subsequent adaptation, could be contributing to off-target antibiotic resistance [21,25].

The goal of this study was to investigate the genetic mechanisms and transcriptome response underlying adaptation to BAC and the effects of BAC adaptation on antibiotic resistance. We used chemical mutagenesis combined with whole-genome sequencing (Mut-seq) to isolate mutants with increased tolerance to BAC and to identify candidate mutations. Additionally, we analyzed the transcriptome of BAC tolerant mutants and the parental strain during exposure to BAC to identify genes and pathways responsive to BAC exposure. Finally, we measured the minimum inhibitory concentration of several antibiotics in the wild-type and mutants to determine if increased tolerance to BAC resulted in changes in antibiotic sensitivity.

## Methods

### Bacterial isolates and culture conditions

Isolate FSL-N1-304 was obtained from the *L. monocytogenes* strain collection at Cornell University’s Food Safety Laboratory. This strain was originally isolated from floor drains in a fish processing facility in New York, USA [26]. All strains were cultured at 37°C in tryptic soy broth media supplemented with 0.6% yeast extract (TSB+YE).

### Chemical mutagenesis using Ethyl Methanesulfonate

Chemical mutagenesis was performed using a previously described protocol with several alterations [27]. Our complete overall experimental design is depicted in Figure 1. First, 1 ml of cells were harvested from an overnight culture through centrifugation at 2,380 x g for 10 minutes. The cells were then washed twice with 1 ml phosphate buffered saline (PBS). After washing, the pellet was resuspended in 120 mM of ethyl methanesulfonate (EMS) suspended in 1 mL of PBS. For a non-mutagenized control, 1 mL of PBS was added to the wild-type (WT) cells. All tubes were incubated for 1 hour at 37°C. Finally, cells were washed three times with PBS to remove EMS and resuspended in 1 mL of PBS.

**Figure 1.**
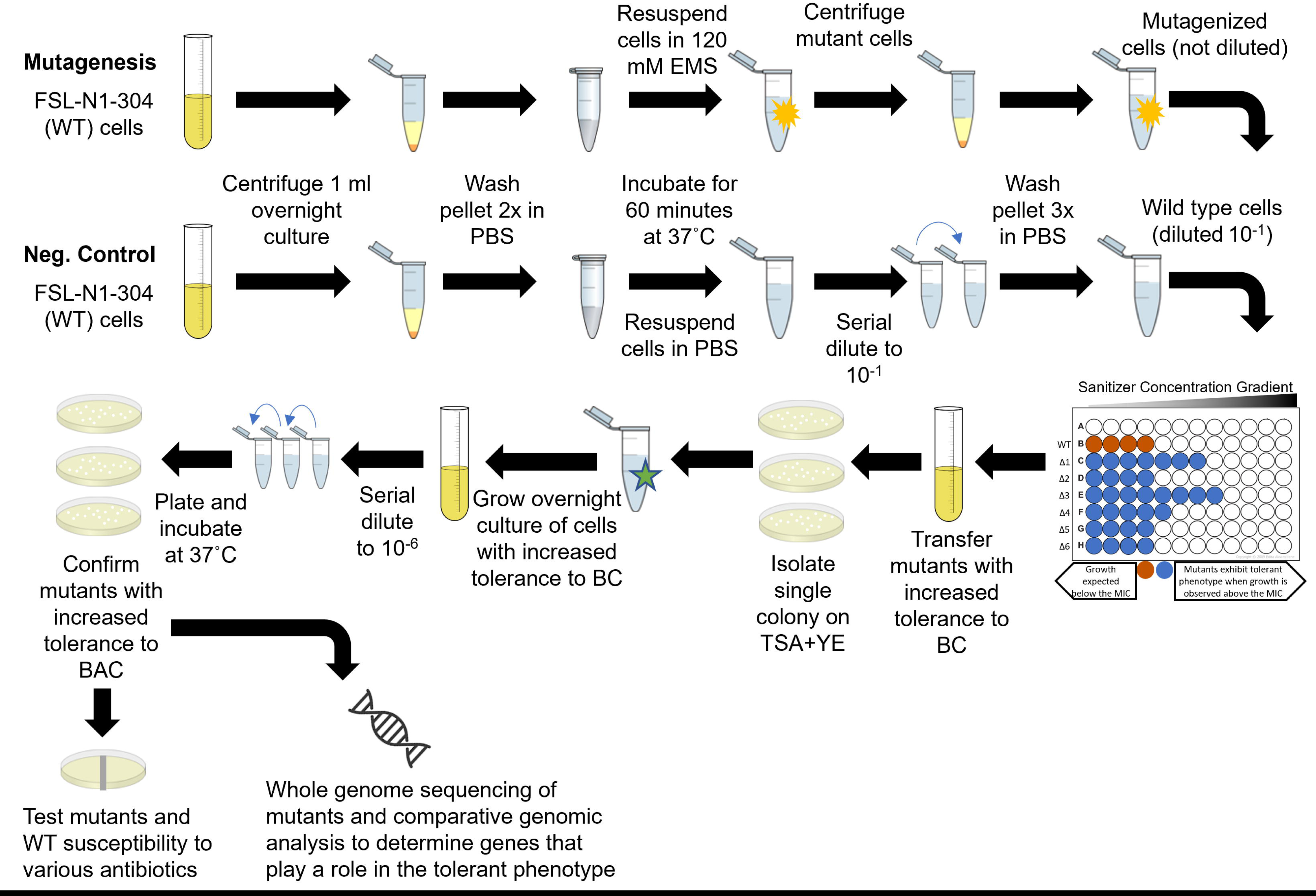
Experimental approach for identifying EMS derived *L. monocytogenes* mutants with increased tolerance to benzalkonium chloride. Experimental approach used for chemical mutagenesis, mutant screening for increased tolerance to benzalkonium chloride, antibiotic susceptibility determination and whole genomic sequencing of mutants with increased tolerance to benzalkonium chloride.

### Determining benzalkonium chloride minimum inhibitory concentration

100 ml of overnight culture of FSL-N1-304 WT cells were inoculated on TS+YE agar plates supplemented with 1, 2, 3, 4, and 5 μg/ml benzalkonium chloride (BAC) and incubated for 48 hours at 37°C to determine the minimum inhibitory concentration (MIC). The MIC was defined as the lowest BAC concentration in which no cell growth was observed. Automated colony quantification and plate imaging was performed using an Interscience Scan1200.

Next, mutants were screened for increased tolerance to BAC in liquid media by inoculating EMS exposed cells into a 96 well plate (BioLite 96 Well Multidish—Thermo Scientific) with BAC concentrations ranging from 0-11 μg/ml, in increments of 1 μg/ml. The top row in each plate represented a negative control which was prepared by adding 50 µl PBS to each well rather than the inoculum. After plating 100 ml of cells treated with EMS following the mutagenesis protocol and 100 ml of untreated cells on TSA+YE plates, we found that exposure to EMS resulted in a 1-log reduction in cell viability. After a 24-hour incubation there was 2.35 x 10^8^ CFU/ml of untreated WT cells, and 8.41 x 10^7^ CFU/ml of EMS treated cell. Thus, to normalize the cell density across WT and mutagenized cells, we diluted the WT cells 10-fold to bring the final cell density of the inoculum to approximately 1 x 10^8^ CFU/ml. The remaining rows were inoculated with 50 µl of the mutant cells, undiluted, immediately after exposure to EMS. The 96 well plate was incubated at 37°C for 24 hours, and optical density at 600 nm (OD_600_) was measured using a BioTek Elx800 microtiter plate reader. Cell density was used to screen the mutants for BAC tolerant phenotypes. We set a threshold for viable cell growth when OD_600_ ≥ 0.1, as previously described [28].

### Validation and MIC determination of increased BAC tolerance in mutants

For mutant cultures that exhibited increased BAC tolerance during screening in 96 well plates, 100 ml was transferred from the wells and inoculated into fresh TSB+YE media and grown overnight. The overnight cultures were then plated on to TSA+YE to isolate single colonies of *mut-1* and *mut-2*. These single colonies were grown overnight in TSB+YE and freezer stocks were prepared in 15% glycerol. All subsequent experimentation was conducted from these stocks.

To validate the BAC tolerant phenotypes observed in the initial screen, overnight cultures of mutants and WT were subjected to the similar BAC screening process as described above using the overnight cultures but in triplicate. After observing that the mutants grew to 6 μg/ml BAC on the 96 well plate during the original screening following mutagenesis, the screening plate was set up similarly, but only included media with 0 - 7 μg/ml BAC, and this time included three biological replicates for the WT and each mutant. After 24 hours OD_600_ readings were collected using a microtiter plate reader. The MIC was recorded as the concentration of BAC in the first well that had an OD_600_ 0.10.

Frozen stocks of the WT and mutants were also used to screen for the MIC of BAC on solid media. Overnight cultures of the WT and mutants were diluted to approximately 1 x 10^4^ CFU/ml and 100 ml of each diluted culture was spread on tryptic soy agar and yeast extract (TSA+YE) plates, as well as TSA+YE plates supplemented with 0, 1, 2, 3, 4, 5, 6, 7 μg/ml BAC. Plates were incubated at 37°C for 48 hours, then images of the plates were obtained and the colonies were quantified using an Interscience Scan1200. The MIC was recorded as the first plate that had no observable colonies for each strain.

### Bacterial growth curve and fitness trade-off analysis

Overnight cultures of WT and mutant cultures were standardized to an OD_600_ of 1.0, and subsequently diluted ten-fold in sterile PBS to achieve a final inoculum of ∼10^4^ CFU/ml. A 5μl volume of this inoculum was then added in triplicate to a 96-well plate (BioLite 96 Well Multidish—Thermo Scientific) with wells each containing 200 ml of TSB+YE (control), TSB+2 μg/ml BAC, or TSB+3 μg/ml BAC. Cultures were then grown for 48 hours at room temperature and growth kinetics were measured using the oCelloScope (Biosense Solutions, Denmark) with the following parameters: auto-illumination, auto-focus, image distance = 4.9mm, images per repetition = 5, repetitions = 48, repetition interval = 1 hour. Total absorption measurements were recorded using a fixed 505nm wavelength and growth kinetic analysis was conducted using the normalized total absorption algorithm.

### *hly* virulence assay

We measured *hly* activity as a proxy for virulence to test if this virulence phenotype was altered in the BAC tolerant mutants compared to the parental strain. *hly* encodes listeriolysin O, which is required for pathogensis. Hemolytic activity was measured for WT and mutant strains based on a previously described method with minor alterations [29]. Briefly, 1 mL of overnight cultures were centrifuged at 14,000 x g for 10 minutes to pellet the bacteria. The supernatant (crude exosubstance) was then transferred to a new microcentrifuge tube and the pellet was discarded. The crude exosubstance was diluted two-fold in PBS in a 96-well plate (ThermoFisher Scientific, Biolite Microwell Plates), leaving a final volume of 100μl in each well. Negative control wells (0% hemolysis) were prepared with 100 μl sterile PBS. Complete hemolysis (100%) control wells were prepared with 100 μl PBS + 1% Triton X-100. A sheep blood solution was prepared by washing pooled, defibrinated sheep blood (Carolina Biological, Burlington, NC) in cold PBS by repeated centrifugation at 600 x g for 10 minutes. Washing steps were repeated until the supernatant appeared clear and colorless. The clean red blood cell (RBC) pellet was then diluted in cold PBS + 20mM cysteine to a 3% RBC suspension. 100 μl of the 3% RBC suspension was then added to each well and mixed by pipetting. The microtiter plate was then incubated at 37°C for 30 minutes. Absorbance readings were performed using an absorbance plate reader (Biotek Instruments Elx800). The following formula was used to quantify the hemolysis percentage: Hemolysis 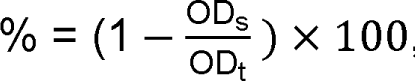, where OD OD_t_ is the difference in optical density at 620nm between the sample and the positive control well and OD_t_ is the difference in optical density at 620nm between the negative and positive control wells.

### Determining the minimum inhibitory concentration of antibiotics

Liofilchem Minimum Inhibitory Test Strips were used to determine the MICs of ciprofloxacin, gentamicin, erythromycin, ampicillin, tetracycline, norfloxacin, penicillin G, and vancomycin in the WT and mutant strains. Overnight cultures of *L. monocytogenes* WT, and mutants were diluted to 10^9^ CFU/ml and spread onto TSA+YE plates using a spreader. Following the manufacturer’s instructions, the Liofilchem strip was positioned on the surface of agar using sterile forceps, and plates were incubated at 37°C for 24 h. Plates were then imaged and MICs were recorded for the WT and mutants for each antibiotic. The MIC was determined by observing where the zone of inhibition ellipse intersected the MIC test strip. This assay was performed in duplicate and averaged to determine the final MIC for each antibiotic. MIC values were classified as susceptible, intermediate, or resistant to the antibiotic according to CLSI breakpoint values for *Staphylococcus spp* [30].

### Whole genome sequencing of FSL-N1-304 and BAC tolerant mutants

From a frozen stock, WT cells were inoculated in TSB+YE media and incubated for 24 hours at 37°C. 1 mL of the overnight culture was centrifuged at 14,000 x g for 10 minutes to pellet the cells. DNA extraction was performed on the pelletized cells using the PureLink Microbiome DNA Purification Kit (Invitrogen; Carlsbad, CA) following the manufacturer’s instructions. For the WT strain, 150-bp paired-end libraries were generated at the Microbial Genome Sequencing Center (Pittsburgh, PA, USA) and sequenced on a NextSeq 2000 sequencer. Low quality reads were trimmed and adapter sequences were removed from Illumina sequences using bcl2fastq version 2.20.0.445 (RRID:SCR 015058) [31]. Oxford Nanopore (ONT) PCR-free ligation library preparation was also used to generate long sequence reads. ONT reads were adapter and quality trimmed using porechop33 version 0.2.3_seqan2.1.1 (RRID:SCR_016967). Hybrid assemblies were generated from Illumina and Oxford Nanopore reads using Unicycler version 0.4.8 and assembly quality was assessed using QUAST version 5.0.2 [32,33]. Gene models were predicted with functional annotations using Prokka version 1.14.5 with default parameters + ‘--rfam’ [34]. Culturing and DNA extraction of mutant strains was performed with the same protocol as the WT and 150-bp paired-end libraries were generated using Illumina sequencing as described above [31–34].

### Identifying mutations in BAC tolerant mutants

TrimGalore v.0.3.7 (RRID:SCR_011847) was used to trim adapter sequences and trim reads at low quality positions with the following parameters: --quality 30, --stringency 1, --gzip, --length 50 [35,36]. All subsequent software was used with default parameters unless otherwise specified. BWA version 0.7.15 (RRID:SCR_010910) was used to map the mutant resequencing data against the ancestral reference genome [37]. Samtools (v 1.4.1) (RRID:SCR_002105) and bamaddrg (https://github.com/ekg/bamaddrg) were used to index and add sample names to the sorted BAM files, respectively [38]. Joint genotyping for SNPs and short indels of the mutants was performed using Freebayes (v 1.3.1) (RRID:SCR_010761) [39]. GATK (v. 4.0.6) was used to convert the resulting VCF files into table format [40]. A SnpEff database was built for the FSL-N1-304 WT reference genome and SNP annotation prediction was performed in the mutant strains using SnpEff (v. 4.1) [41]. Large-scale Blast Score Ratio (LS-BSR) was used to investigate gene presence/absence mutations using default parameters with the exception of “-b blastn -c cdhit” [42]. The InterPro server was used to predict protein domains in candidate proteins (https://www.ebi.ac.uk/interpro/) [43].

### RNA-sequencing and differential gene expression analysis

We performed RNA-seq on the WT and mutant strains in the presence of BAC in order to compare their transcriptome profiles. Each culture was revived from freezer stocks by growing in TSB+YE overnight at 37C. 100μl of overnight cultures were then passaged into TSB+YE containing 2 μg/ml BAC and grown to mid-log phase (10 hours). Bacteria were then centrifuged at 14,000 x g for 10 minutes to obtain bacterial pellets which were rapidly frozen using dry ice. RNA extraction, ribosomal depletion library preparation and sequencing were performed at SeqCenter (Pittsburg, PA). Sequencing was performed on a NovaSeq 6000, generating 2×51bp reads. Three biological replicates were sequenced for each sample. Adapter trimming and demultiplexing were performed with bcl-convert (v4.0.3) [31]. Genome indexing and mapping were performed using bwa (v0.7.17) with default parameters [37]. Samtools (v1.14) was used to sort and index BAM files [38]. Bedtools2 (v2.30.0) was used to extract read counts per transcript from gene coordinates in the sorted BAM files [44]. The DESeq2 (v.1.38.3) package was used in R (version 4.2.3 and RStudio 2023.03.1+446) to normalize read counts and identify differentially expressed genes [45]. Differential expression thresholds were defined using log-fold change [log2(FC)] values >1 (upregulated) and <-1 (downregulated). A p-value significance threshold was also defined using the multiple test corrected p-value 1.6E-5. Principal component analysis (PCA) was performed on DESeq2 normalized read counts using JMP Pro (version 17) to examine the relationship between samples and replicates [46]. Volcano plots of gene expression (DESeq2 normalized read counts) and p-value were generated using ggplot2 (v.3.4.1) [47]. KEGG annotations were generated for the whole genome and used to identify statistically significant overrepresentations of specific KEGG terms among differentially expressed genes using Fisher’s exact tests in *R* using an FDR cutoff of 0.01 [48].

## Results

### BAC MIC in FSL-N1-304 and Mutants

The BAC MIC for FSL-N1-304 was measured in both liquid culture and solid media to establish a baseline MIC for all subsequent screening steps. All experiments were performed in triplicate and averaged to calculate the baseline MIC. In the liquid culture assay, the BAC MIC for the WT strain was 4 μg/ml (Figure 2a). For the solid media assay, the BAC MIC was 3 μg/ml (Figure. 2b). During our mutant screening, we isolated two mutants with increased tolerance to BAC (*mut-1* and *mut-2*). *mut-1* and *mut-2* displayed BAC MICs of 7 μg/ml in liquid culture, and 4 and 5 μg/ml in solid culture, respectively (Figure 2a-b). We also measured the growth kinetics of the WT, mut-1, and mut-2 in the presence of 0, 2, and 3ug/ml BAC in liquid TSB+YE media over the course of 48 hours. WT, *mut-1* and *mut-2* had nearly identical growth patterns in TSB+YE (Figure 3), suggesting there is not a growth tradeoff in the absence of BAC in the mutant strains. Additionally, growth of the WT diminished compared to the mutants as the BAC concentration was increased (Figure 3).

**Figure 2.**
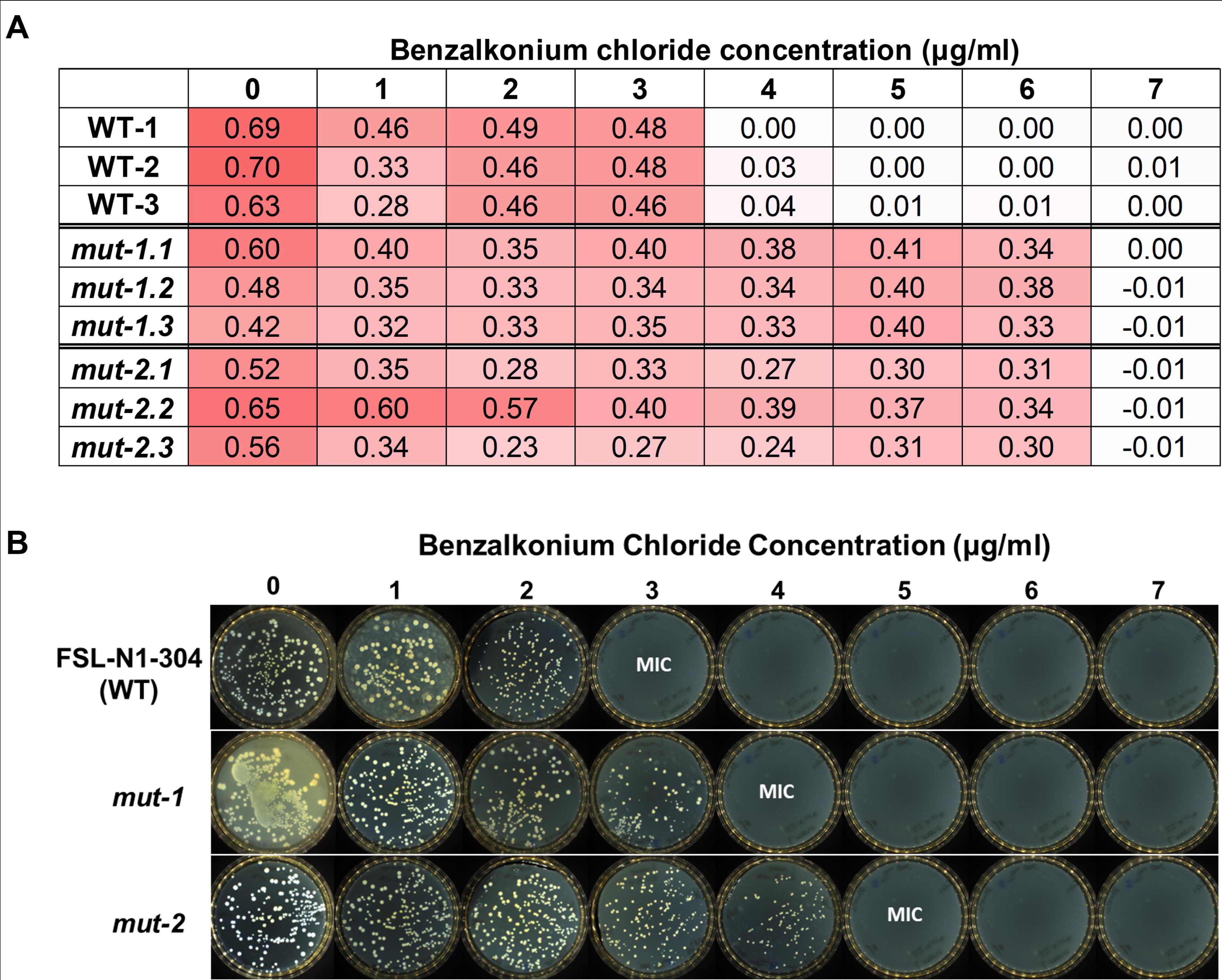
Minimum inhibitory concentration of benzalkonium chloride in WT and mutant *L. monocytogenes* strains. **A)** Columns represent samples (WT, *mut-1* and *mut-2*) and biological replicates, whole rows show the concentration (ug/ml) of benzalkonium chloride. Each component of the matrix shows the OD value after overnight growth. Growth was considered entirely inhibited when OD values < 0.1. The red shading represents OD value, with light and dark red representing low and high OD values. **B)** Minimum inhibitory concentrations of *L. monocytogenes* WT and mutants to benzalkonium chloride on solid media. 100 μl of overnight culture from the WT, mut-1 and mut-2 were spread onto TSA+YE plates with 0, 1, 2, 3, 4, 5, 6, and 7 μg/ml benzalkonium chloride and incubated at 37°C for 24 hours. Plates were then imaged on an Interscience Scan1200. Benzalkonium chloride minimum inhibitory concentration (MIC) was determined as the concentration at which no colonies were visible.

**Figure 3.**
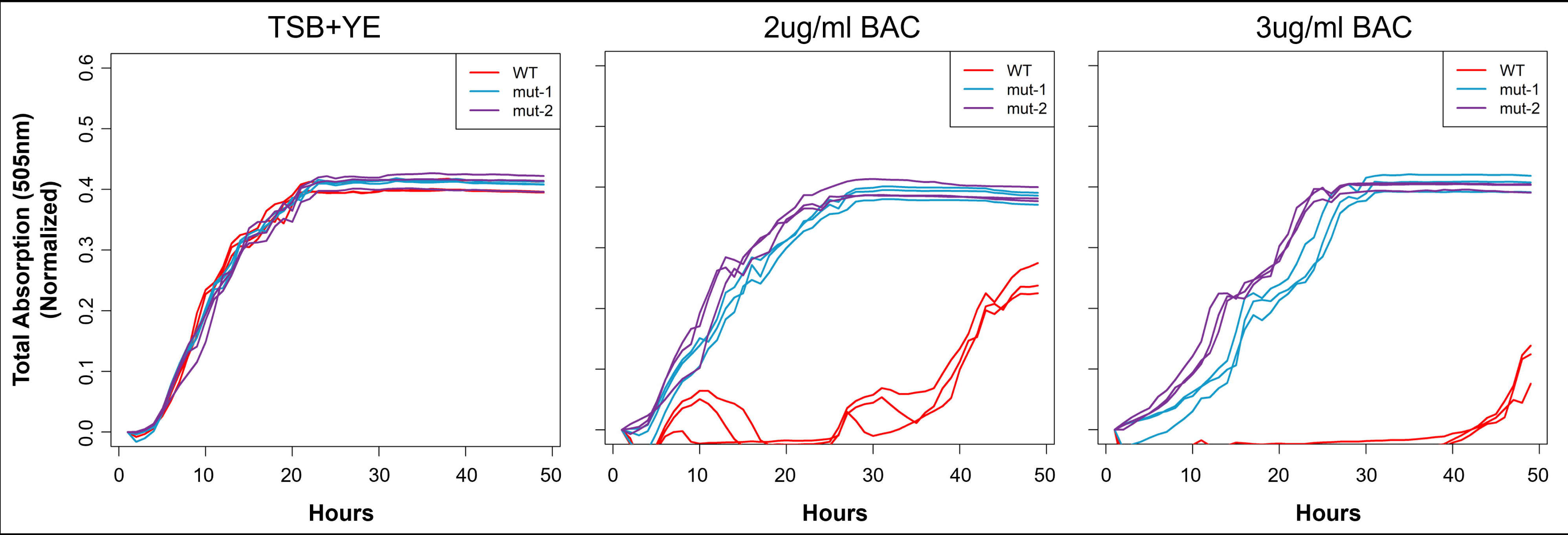
Growth rate analysis of WT, *mut-1*, and *mut-2* in response to increasing concentrations of benzalkonium chloride. Each culture was grown at room temperature for 48 and the growth rate was measured using the oCelloScope. (A) Growth curve of WT and mutant strains based on the total absorption (505nm wavelength) values in each microtiter well. (B) Growth curve of WT and mutant strains based on background corrected absorption (BCA) values. Growth rate was measured in 3 biological replicates for each sample. WT, *mut-1*, and *mut-2* are depicted by red, light blue and purple lines respectively.

### BAC tolerant mutants show increased tolerance to some antibiotics

Previous studies show that BAC tolerant *L. monocytogenes* isolates often also show increased tolerance to some antibiotics [19,21]. To test whether this phenomenon was observed in our BAC tolerant mutants, we measured the MIC of ciprofloxacin, gentamicin, erythromycin, ampicillin, tetracycline, norfloxacin, penicillin G, and vancomycin in the WT and mutant strains (Figure 4, Table 1). The MICs for erythromycin, ampicillin, tetracycline and penicillin G in the WT and mutants were either identical, decreased in the mutants, or showed a minor increase in the mutants but would still be considered “sensitive” to the antibiotic based on *Staphylococcus spp.* MIC breakpoints [30]. Additionally, we observed a near doubling of vancomycin MIC from 0.44 μg/ml in the WT to 0.75 and 0.875 μg/ml in in *mut-1* and *mut-2*, respectively, and an increase in gentamicin MIC from 1.25 μg/ml to 4 μg/ml in *mut-2*, respectively (Figure 4a, Table 1). The mutant vancomycin and gentamicin MICs are still considered “sensitive” to the antibiotics, based on MIC breakpoints for other species. For ciprofloxacin, we observed an increase in MIC from 0.5 μg/ml in the WT to 2 μg/ml in *mut-1*, which is considered “intermediate” tolerance (Table 1, Figure 4b). Lastly, we observed a substantial increase in norfloxacin MIC from 2.5 μg/ml in the WT, to 16 and 8 μg/ml in *mut-1* and *mut-2*, respectively (Table 1, Figure 4b). Norfloxacin MICs of 8 and 16 μg/ml are considered “intermediate” and “resistant”, respectively.

**Figure 4.**
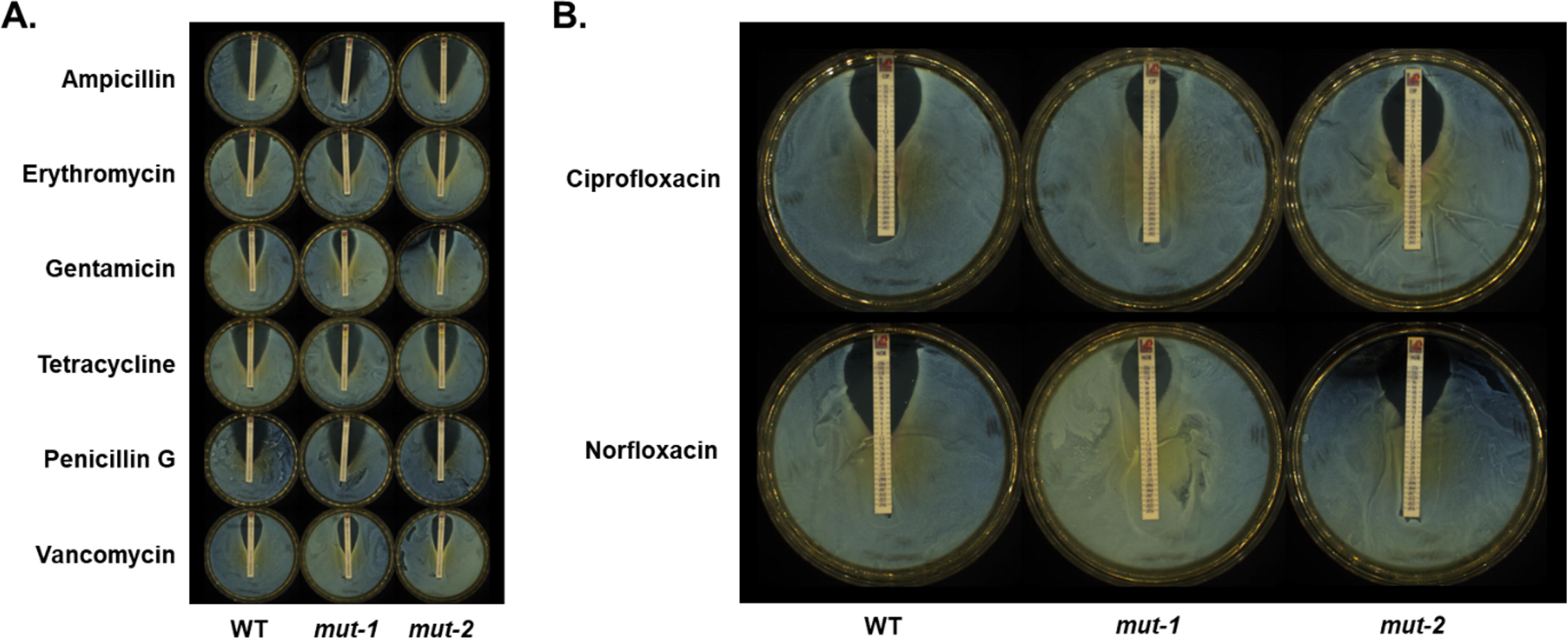
Minimum inhibitory concentrations of various antibiotics in the wild type and mutants with increased tolerance to benzalkonium chloride. The minimum inhibitory concentrations of ampicillin, erythromycin, gentamicin, tetracycline, penicillin G, and vancomycin (A) and the fluoroquinolones, ciprofloxacin and norfloxacin, (B) were measured in the WT, *mut-1* and *mut-2* using Liofilchem Minimum Inhibitory Test Strips. 10^9^ CFU/ml cells were spread onto the TSA-YE plates and incubated at 37°C for 24 hours to create a lawn. The minimum inhibitory concentration was determined as the point where the edge of the inhibition ellipse intersected with the MIC Test Strip.

**Table 1.**

| Antibiotic | MIC for WT (µg/ml) | mut-1 MIC (µg/ml) | mut-2 MIC (µg/ml) |
| --- | --- | --- | --- |
| Ampicillin | 0.0555 | 0.016 | 0.0555 |
| Erythromycin | 0.315 | 0.315 | 0.22 |
| Gentamicin | 1.25 | 1.5 | 4* |
| Penicillin G | 0.035 | 0.035 | 0.0275 |
| Tetracycline | 0.157 | 0.1875 | 0.1875 |
| Vancomycin | 0.44 | 0.75 | 0.875* |
| Ciprofloxacin | 0.5 | 2* (I) | 0.5 |
| Norfloxacin | 2.5 | 16** (R) | 8* (I) |

### Assessing virulence potential through hemolytic assay

To test whether the BAC tolerant phenotype is linked with *L. monocytogenes* virulence, we measured *hly* activity, an essential component of virulence, in the WT and mutant strains. *hly* activity was measured as the percent of hemolysis in goat blood. The parental strain had an average hemolysis of 6.05%, while *mut-1* and *mut-2* had significantly higher hemolysis activities of 67.71% (p-value = 0.00093) and 45.45% (p-value = 0.012), respectively (Figure 5).

**Figure 5.**
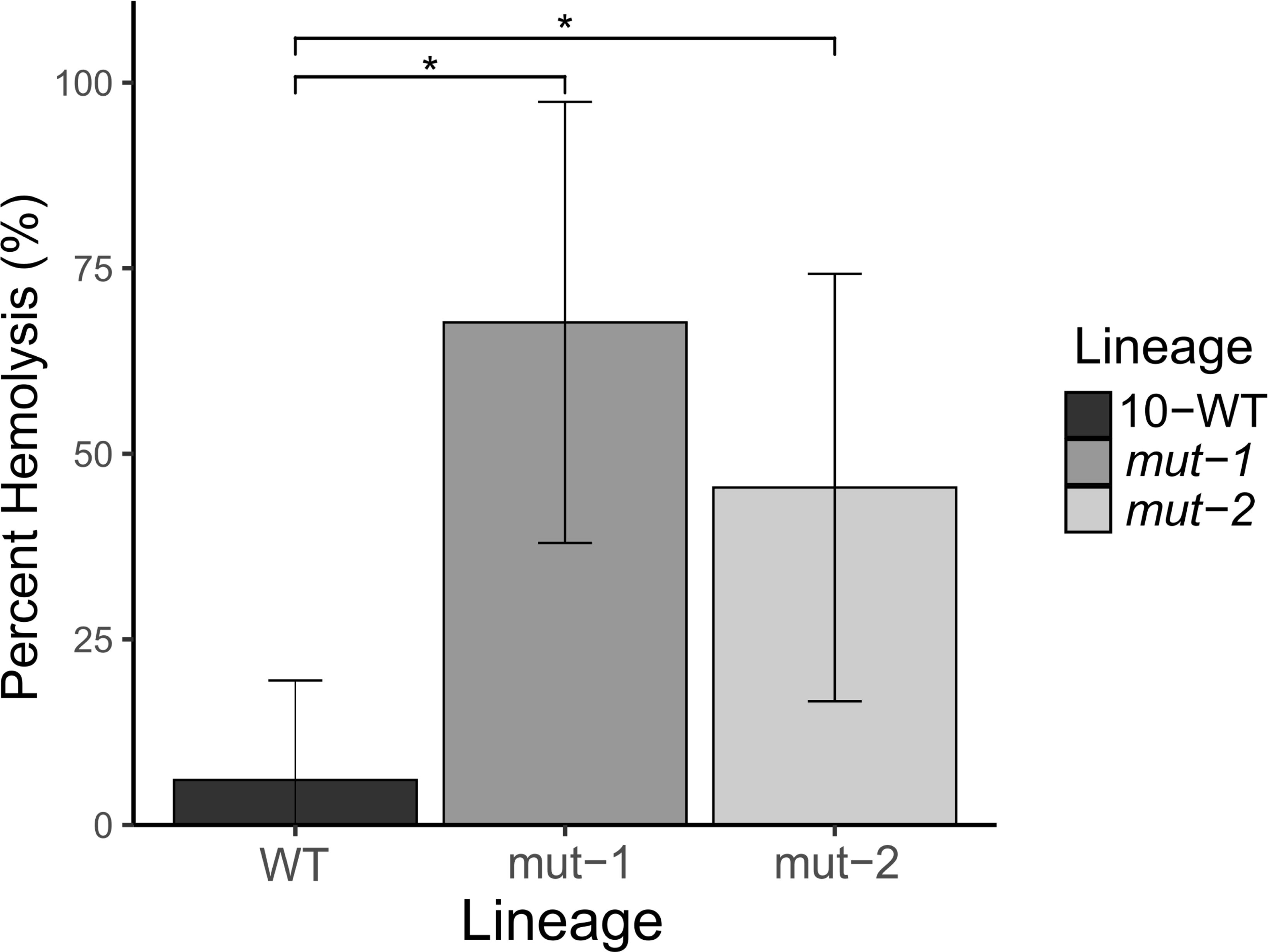
Hemolytic activity is significantly increased in BAC-tolerant mutants. To investigate the relationship between BAC tolerance and virulence, we conducted a hemolysis assay using *mut-1*, *mut-2*, and the parental strain. Percent hemolysis, which is a semi-quantitative proxy for the expression of key *L. monocytogenes* virulence factor, hly, is reported. Error bars represent the standard deviation of percent hemolysis for each lineage. Statistically significant values between WT, *mut-1* (p-value = 0.00093), and *mut-2* (p-value = 0.012) are indicated by an asterisk.

### Genomic analysis of mutants

To investigate the EMS induced mutations that may account for the increased tolerance to BAC and antibiotics, we generated a complete genome assembly for FSL-N1-304 using Illumina and Oxford Nanopore Technologies sequencing and predicted gene models and functional annotations (NCBI SRA run accession numbers SRR17006605 and SRR17067427 for the Illumina and ONT data, respectively). The genome was assembled into two contigs consisting of the genome (∼3 Mb) and a plasmid (pLIS10—63.8Kbp) with a total of 3,033 protein coding gene models (NCBI accession number GCA_021391455.1). This high-quality genome was used as a reference to map Illumina whole-genome resequencing data for *mut-1* and *mut-2*. We identified 3 and 29 mutations in the *mut-1* and *mut-2* genomes, respectively. For *mut-1*, consistent with EMS resulting mainly in G:C>A:T transitions, we observed three G>A missense mutations in genes with predicted functions as a diacylglycerol kinase, a permease, and a resuscitation-promoting factor (Table 2; Figure 6). In *mut-2*, we observed seven C>T transitions, 20 G>A transitions, one G>T transversion, and one single nucleotide deletion (Table 2). Twenty two of the 29 mutations in *mut-2* were missense mutations, three resulted in premature stop codon mutations, and a single nucleotide deletion resulted in a frameshift mutation (Table 2). Interestingly, none of the genes with mutations in *mut-1* and *mut-2* overlapped, suggesting different mechanisms of increased BAC tolerance.

**Figure 6.**
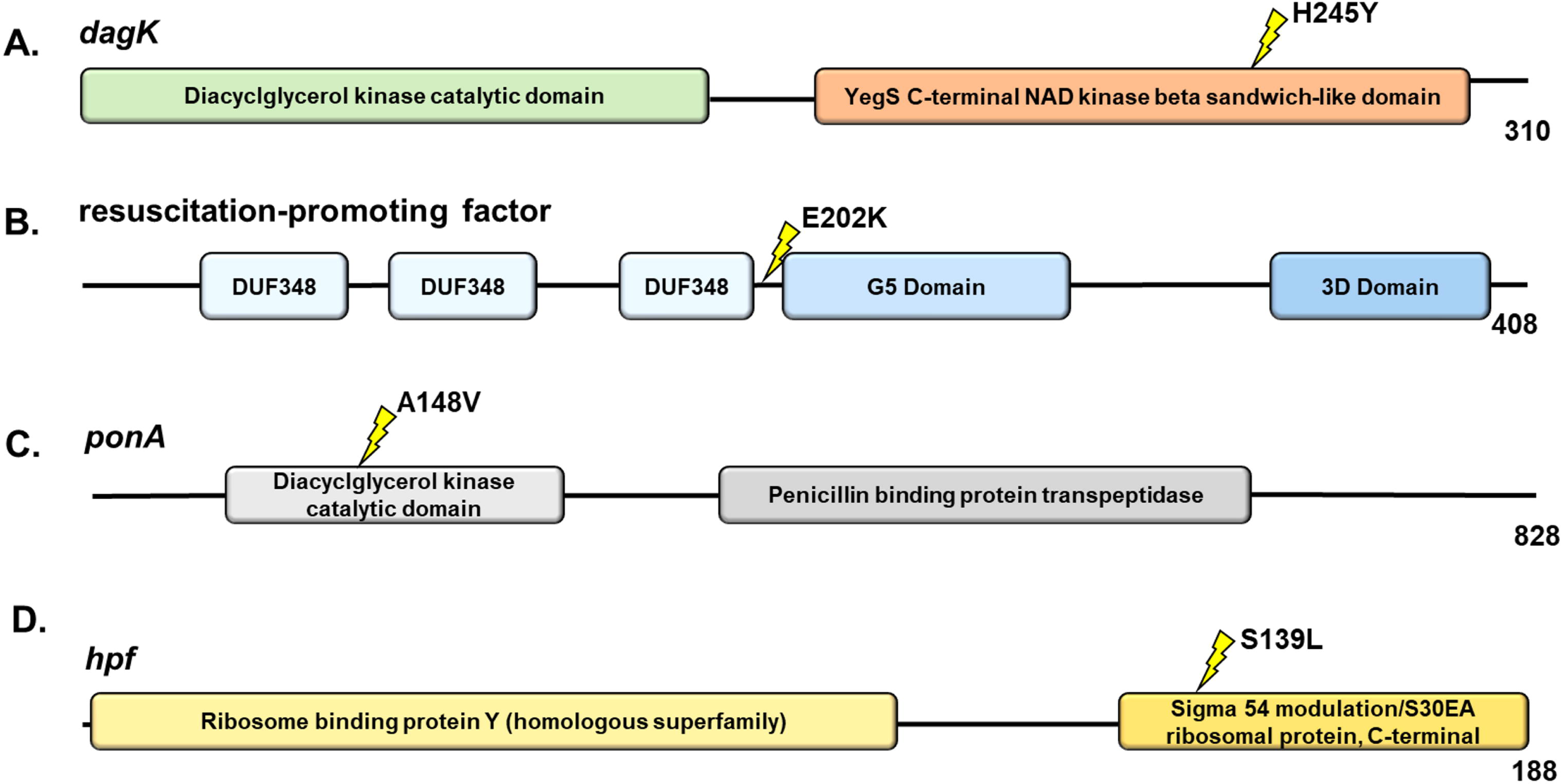
Candidate proteins in mutants with increased resistance to benzalkonium chloride. The common *L. monocytogenes* gene name is provided for each protein. InterPro functional domains are represented by colored ellipses. The yellow lightning bolt symbol represents the site of the nonsynonymous mutations. (A and B) *dagK* and resuscitation-promoting factor mutations were present in *mut-1*, while (C and D) *ponA* and *hpf* mutations were present in *mut-2*. The number at the end of the protein represents the length of the protein in amino acids.

**Table 2.**
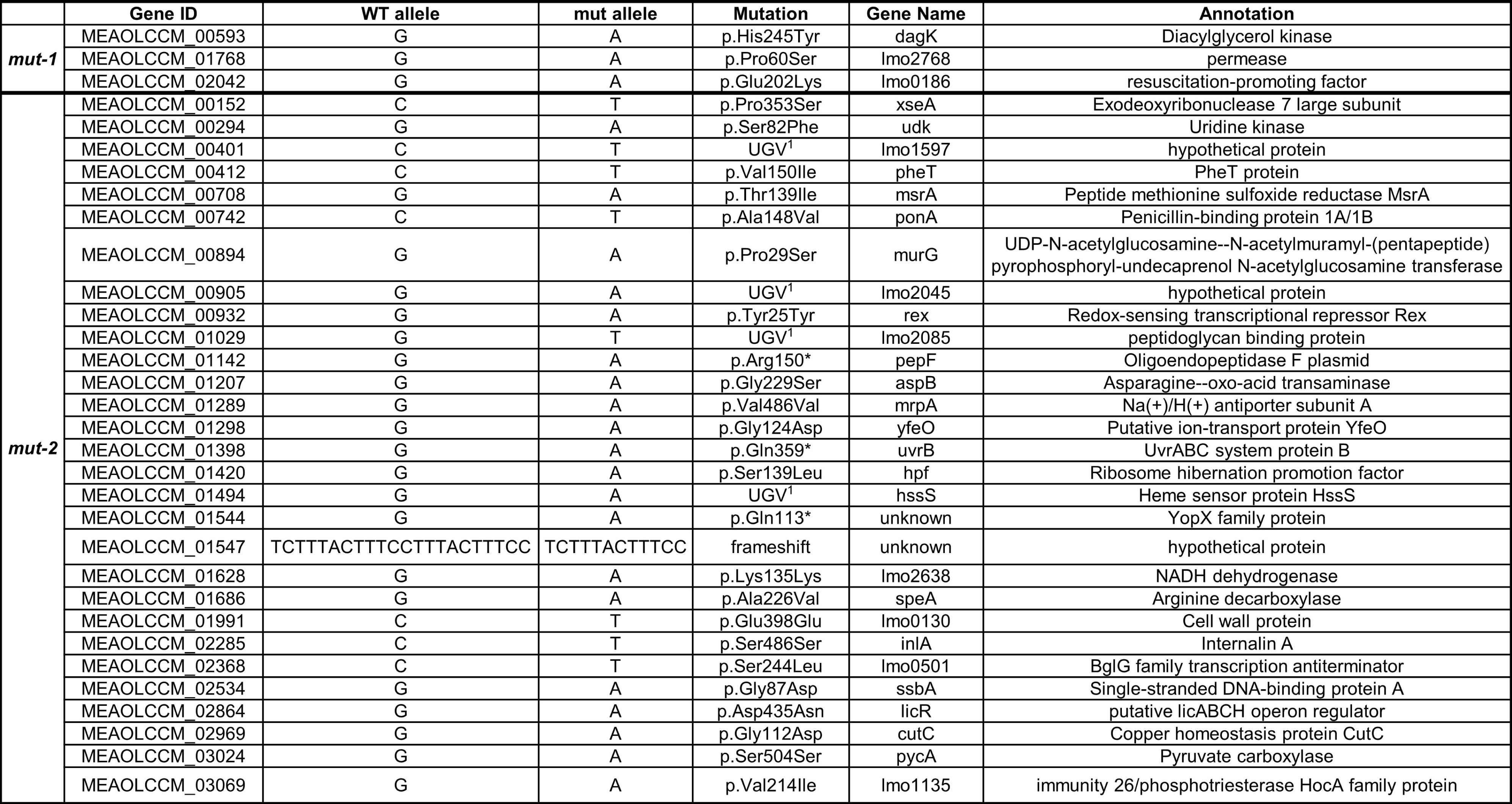

Additionally, we investigated whether gene gain or loss in the mutants may have changed after mutagenesis using the LS-BSR pipeline [42]. Consistent with predictions from EMS mutagenesis, we did not detect any gene gains or losses in *mut-1* or *mut-2*, suggesting that structural variants did not influence the increase in BAC tolerance.

### Differential gene expression analysis between WT and BAC-tolerant mutants

We performed RNA-sequencing of the WT, *mut-1* and *mut-2* in biological triplicates during growth in TSB+YE containing 2 μg/ml BAC for 10 hours (mid-log phase) to examine transcriptome profiles differences across samples. We performed principal components analysis (PCA) of DESeq2 normalized read counts (Figure 7c). Biological replicates for each sample clustered together, suggesting the biological replicates had similar transcriptome profiles, and that there are major differences in the transcriptome response to BAC across samples. We independently identified differentially expressed genes in *mut-1* and *mut-2* relative to the WT. We identified 282 up-regulated genes and 134 down-regulated genes in *mut-1*, and 136 up-regulated and 45 down-regulated genes in mut-2 (Figure 7). *mut-1* and *mut-2* share 55 up-regulated genes and 15 down-regulated genes (Supplementary Table 2a-b). Genes of the phosphotransferase (PTS) system (including: *bglG, dhaL, dhaM, ulaA, fruA, manZ, licABC*) were up-regulated in both *mut-1* and *mut-2* compared to the WT. KEGG pathway enrichment analysis supports this observation, revealing that KEGG terms corresponding to the PTS system were overrepresented amongst all upregulated genes in both *mut-1* (p-value = 0.000452) and *mut-2* (p-value = 0.00112). Additionally, two internalin-encoding genes, *inlB* and *inlH*, the *groEL-groES* chaperonin system, transcriptional antiterminator *lmo0425*, and multidrug transporter *lmo2463* were all upregulated in both *mut-1* and *mut-2*. Interestingly, KEGG pathway enrichment analysis also revealed that genes involved in amino acid biosynthesis were overrepresented in the up-regulated genes of *mut-1*, while amino acid metabolism was overrepresented in up-regulated genes of *mut-2* (Table 3).

**Figure 7.**
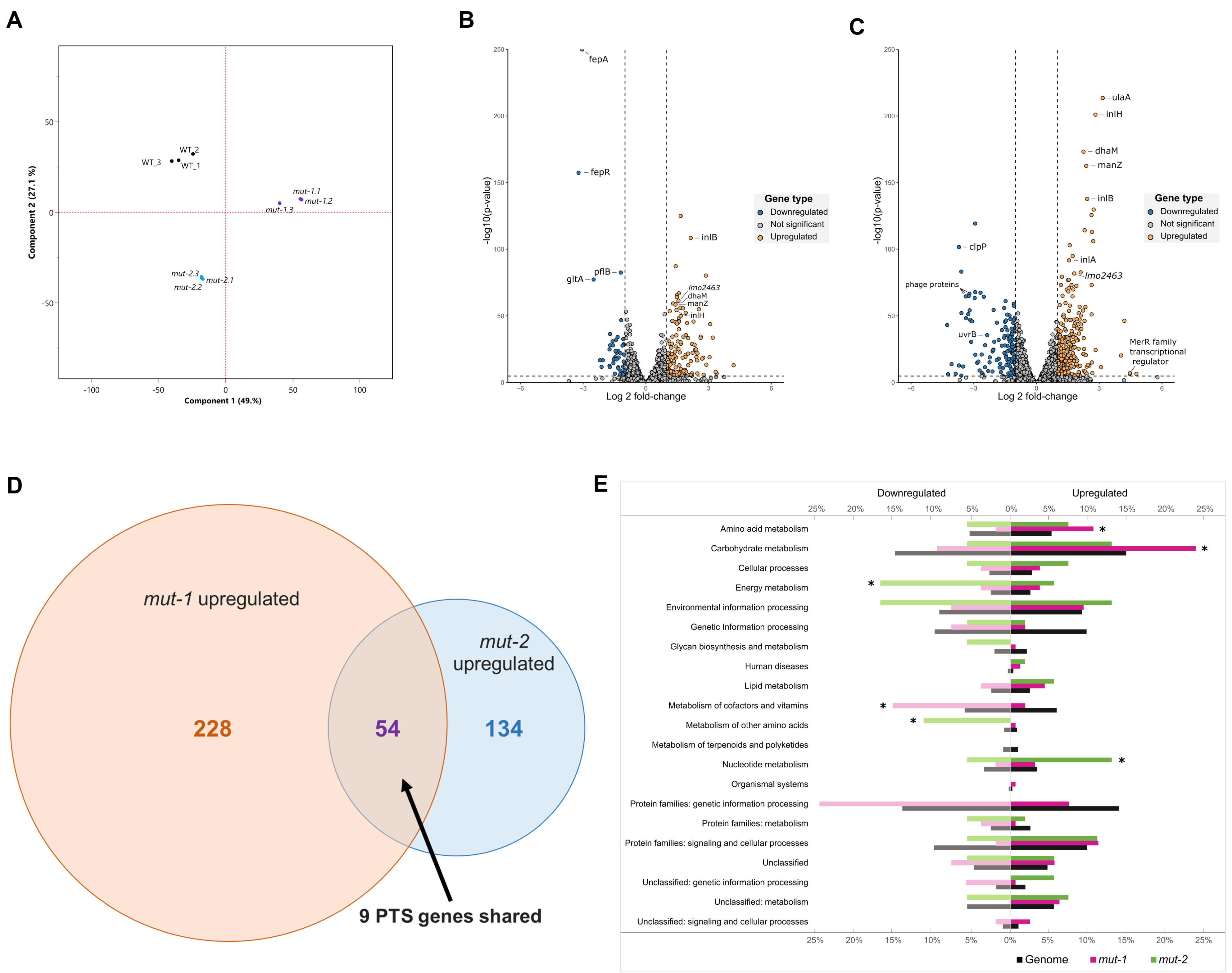
Transcriptome response to BAC stress in WT and mutants. (A) Principal component analysis of RNA-sequencing data reveals distinct clustering of WT and mutant strains. PCA was conducted on genome-wide normalized DESeq2 read counts. *mut-1* and *mut-2* replicates are displayed using purple dots and light blue dots, respectively. WT replicates are represented with black dots. (B) Volcano plots displaying differential gene expression analysis results for *mut-1* and (C) *mut-2* show differential gene expression of mutants relative to the parental strain. Vertical dashed lines represent differential expression thresholds, which were defined using log-fold change [log2(FC)] values >1 (upregulated) and <-1 (downregulated). The horizontal dashed line represents the significance threshold defined using a multiple test corrected p-value: (significance level)/(total # of genes)= 0.05/3201 = 1.6E-5. (D) Venn diagram comparison of all upregulated genes identified in *mut-1* and *mut-2*. The PTS system accounted for 9 out of 52 (16.67%) shared upregulated genes. (E) Enrichment of KEGG terms among differentially expressed genes. The proportion of genes belonging to each functional category within (1) the total genome, (2) differentially expressed genes in mut-1, and (3) differentially expressed genes in mut-2 correspond to black, pink, and green bars, respectively. Asterisks indicate significant overrepresentations (p-value <0.01).

We identified *fepR,* the negative regulator of the efflux pump *fepA*, in the subset of genes down-regulated in both *mut-1* and *mut-2* (Figure 7a-b). Interestingly, despite the downregulation of *fepR*, we found that *fepA* was highly downregulated (LFC = -3.05, *p-value* = 0) in *mut-2*. We also observed transcriptome profile differences between the two mutants in the *ilv-leu* operon, which is highly up-regulated only in *mut-1* (*leuA*—LFC = 2.35, p-value = 2.91E-08; *leuB*—LFC = 2.52, p-value = 2.15E-13; *leuC*—LFC = 2.40, p-value = 1.28E-12; *leuD*—LFC = 2.46, p-value = 7.07E-14; *ilvA*—LFC = 2.26, p-value = 5.63E-34; *ilvB*—LFC = 1.82, p-value = 3.32E-06; *ilvC*—LFC = 2.27, p-value = 2.12E-09; *ilvH*—LFC = 2.02, p-value = 1.76E-08). The *ilv-leu* operon is involved in branched-chain amino acid synthesis and has been associated with *L. monocytogenes* survival under harsh, host-associated conditions [49,50]. Only four of the candidate genes with mutations in either *mut-1* or *mut-2* were differentially expressed *(mut-1*: *lmo0186* and mut-2: *rex, yfeO, hpf*).

## Discussion

Here, we used EMS mutagenesis to generate *L. monocytogenes* mutants with increased tolerance to BAC in an effort to understand the potential genetic underpinnings and transcriptome response to BAC tolerance, and the off-target effects of BAC tolerance on antibiotic sensitivity. We conducted this study with *L. monocytogenes* FSL-N1-304 because this strain lacks genomic elements, such as *bcrABC*, *qacH* and *Tn6188*, which are involved in QAC resistance [51–53]. For instance, analysis of over 116 *L. monocytogenes* isolates from diverse serotypes and origins, revealed that 71 of the 72 strains displaying BAC resistance harbored *bcrABC* [51], while another study showed that strains harboring *Tn6188* have BAC MICs twice as high as strains lacking *Tn6188* [53]. Thus, the isolation of FSL-N1-304 mutants displaying increased BAC tolerance could reveal novel genetic elements and mechanisms involved in BAC tolerance.

We generated two mutants with elevated BAC MICs in both liquid and solid media compared to the parental FSL-N1-304 strain (Figures 2 and 3). Additionally, *mut-1* and *mut-2* harbored resistance and increased tolerance to norfloxacin, respectively, while the parental FSL-N1-304 strain was sensitive to the antibiotic (Table 1 and Figure 4B). We sequenced the genomes of the WT, *mut-1* and *mut-2* to identify candidate mutations involved in increased BAC tolerance. Interestingly, the number of mutations we observed between our two mutants differed by an order of magnitude (Table 2). In two recent EMS mut-seq studies in *L. monocytogenes,* 1 to 6 mutations were observed per mutant [54,55]. *mut-2* contained 29 mutations, and the majority of these mutations were in G:C > A:T transitions, consistent with expectations from EMS mutagenesis. However, we did observe a mutation in *uvrB* (which encodes the nucleotide excision repair enzyme UvrABC system protein B) that results in a premature stop codon and truncated protein that lacks the C-terminus helicase C, UvrB_YAD/RRR_dom, and UVR_dom domains (Table 2). This mutation is likely responsible for the higher number of mutations observed in *mut-2*. *mut-1* contained only three nonsynonymous mutations that differentiate it from the parental FSL-N1-304 strain. One of these mutations occurred in *dagK*, a gene encoding a diacylglycerol kinase (Figure 6a). In *Escherichia coli*, DAGK recycles diacylglycerol that is produced by membrane-derived oligosaccharide biosynthesis [56]. Interestingly, in *Acinetobacter baumannii*, *dagK* was significantly up-regulated in a colistin-resistant strain compared to its colistin-susceptible parental strain [57]. Both colistin and BAC disrupt the membrane, and alterations to expression or structure may improve cell membrane integrity, underlying a potential mechanism for increased BAC tolerance in *mut-1*. Additionally, *mut-1* contained a nonsynonymous mutation in a predicted permease encoding gene (Table 2). The P60S mutation is predicted to occur outside the membrane in the extracellular region. Lastly, we observed a mutation in a gene encoding a predicted resuscitation-promoting factor that is also annotated in the Gene Ontology category peptidoglycan turnover (GO:0009254) (Figure 6b). As a gram-positive bacterium, the *L. monocytogenes* cell wall is composed of a rigid layer of peptidoglycan that acts as a barrier to the external environment. It has previously been hypothesized that modifications to peptidoglycan synthesis could alter membrane fluidity and could be involved in BAC resistance [18].

Though linking mutations to the increased BAC tolerance phenotype in *mut-2* is more challenging because of the relatively large number of mutations observed, we did identify several interesting candidates. *mut-2* contained a nonsynonymous mutation (A148V) in the penicillin-binding protein (PBP) encoding gene, *ponA* (Figure 6c). This protein is involved in polymerizing and modifying peptidoglycan, making it an interesting candidate for the BAC tolerance phenotype. While PBPs have not been implicated in QAC resistance, their presence has been associated with *L. monocytogenes* resistance to β-lactam antibiotics [58]. Additionally, we observed a nonsynonymous mutation (S139L) in the gene encoding for the ribosome hibernation promotion factor (HPF) (Figure 6d). In *L. monocytogenes*, *hpf* knockout mutants showed greater uptake of gentamicin and generally, were more sensitive to aminoglycosides during stationary phase [59]. This mutation is an interesting candidate, as we observed an increase in gentamicin MIC from 1.25 μg/ml in the parental strain to 4 μg/ml in *mut-2* (Figure 4a, Table 1). Additionally, *L. monocytogenes* strains harboring plasmid pLMST6, which contains *emrC*, a multidrug efflux pump transporter, showed increased tolerance to both BAC and gentamicin [60], suggesting increased tolerance to these compounds may achieved through similar paths.

We hypothesized that there are two separate efflux mechanisms acting as contributors to the decreased antimicrobial susceptibility observed in these mutants because the transcriptome profiles of the two mutants reveal differential expression of *fepR* and *lmo2463*. *fepR* encodes a transcriptional regulator that negatively regulates its downstream target, FepA, a multi-drug efflux pump that is linked to both QAC and fluoroquinolone resistance [21,22,61]. One study showed that a loss-of-function frameshift mutation in *fepR* resulted in up-regulation of *fepA* and subsequent resistance to norfloxacin and ciprofloxacin [24]. Another study experimentally evolved *L. monocytogenes* for increased tolerance to BAC and didecyldimethyl ammonium chloride and showed that these adapted lineages were also resistant to ciprofloxacin stemming mainly from missense and nonsense mutations in *fepR* [22]. Interestingly, the increased tolerance to ciprofloxacin was only observed in *mut-1* and was not as drastic as the increased MIC for norfloxacin (Figure 4b). We did not observe mutations in *fepR* or *fepA* in the mutants, which initially suggested that the increased tolerance to BAC and fluoroquinolones occurred through pathways independent of the *fepR/fepA* genotype. However, our RNA-seq results suggest that the *fepR/fepA* mechanism may be still contributing to the BAC mutant phenotypes. *fepR* was downregulated in both mutants in the presence of BAC, albeit the relative expression was considerably lower in *mut-2* compared to *mut-1* (*mut-1:* log2[FC] = -3.22 and *mut-2*: log2[FC] = -1.03). Surprisingly, *fepA* was significantly down-regulated in *mut-1*, and not significantly differentially expressed in *mut-2* (Figure 7). This is the opposite result that we would expect considering that *fepR* negatively regulates *fepA,* as well as the fact that a higher MIC observed in *mut-2* (Figure 2) compared to *mut-1*.

The second efflux pump mechanism that we hypothesize may to be contributing to decreased antimicrobial susceptibility is the MmpL family transporter protein encoded by *lmo2463*. MmpL family transporters are a class of multi-drug resistance proteins that increase resistance to fluoroquinolones in *E. coli* [62]. Other studies have shown that MmpL family proteins were induced under acid stress in *L. monocytogenes* [63]. We found that *lmo2463* was upregulated in both mutants when grown in the presence of BAC stress. Additionally, considering the increased tolerance to fluoroquinolone antibiotics compared to the WT, the MmpL transporter is a potential candidate of BAC and fluoroquinolone tolerance. Efflux pump deletion mutants will need to be generated in future work to determine if synergistic effects exist between *fepR/fepA* and the MmpL efflux mechanism.

Collectively, our results offer potential insight to the aforementioned hypothesis that variability in peptidoglycan synthesis contributes to BAC tolerance as we observed a significant up-regulation of genes of the phosphotransferase system (PTS), which is involved in cell wall turnover and peptidoglycan modulation [18,64]. Enrichment analysis of KEGG pathways also revealed that *mut-1* and *mut-2* contained 20 and 14 up-regulated PTS genes, respectively. The PTS system in *L. monocytogenes* (as well as other bacterial pathogens) is associated with tolerance mechanisms to various stressors including sanitizers and antibiotics [63,65–69].

Several PTS genes that were up-regulated in *mut-1* and *mut-2* are highlighted in Figure 7a-b, including *ulaA, dhaM*, and *manZ*. Another study on benzalkonium chloride tolerance in *L. monocytogenes* found that *manZ* mutations were only observed in benzalkonium-adapted strains, which provides further support that the PTS system is heavily involved in biocide tolerance [22]. Further, an additional study demonstrated similar findings, observing that that PTS genes were highly upregulated in *L. monocytogenes* when exposed to benzethonium chloride (BZT), another QAC [65]. The PTS system has other implications for stress tolerance as demonstrated by the disruption of the mannose-specific PTS operon *mpt* and overexpression of the *bgl* operon, which caused increased resistance to bacteriocins [67,68,70].

As shown in Figure 5, a significant increase in hemolytic activity was observed in both *mut-1* (p-value = 0.000931) and *mut-2* (p-value = 0.0125) compared to the WT, suggesting that increased BAC tolerance may be positively correlated to higher hemolytic activity. Additionally, several internalin-encoding genes involved in virulence, including *inlB* and *inlH*, were upregulated in both *mut-1* (*inlB*—LFC = 2.42, p-value = 1.84E-138; *inlH—*LFC *=* 2.82, p-value = 1.01E-201) and *mut-2* (*inlB*—LFC = 2.15, p-value = 3.61E-109; *inlH—*LFC *=* 1.46, p-value = 4.25E-55) (Figure 7a-b). Expression of *InlH* has been previously shown to be associated with *L. monocytogenes* stress response, as it is under σ^B^-dependent regulation [71]. Transcriptional analysis showed no significant difference in the expression of *hly* or other key virulence factors such as *prfa, actAB,* or *plcAB* under BAC stress. We did, however, observe significant up-regulation of the ATPase-encoding gene, *clpC* (*mut-1*: LFC = 1.58, p-value = 9.92E-104 and *mut-2*: LFC = 1.14, p-value = 3.79E-54), which has been previously associated with increased virulence and increased susceptibility to a variety of stressors [72,73]. In *mut-1*, we also observed upregulation of the *ilv-leu* operon, which is primarily associated with branched-chain amino acid synthesis but has also been shown to be under nutrient-dependent negative regulation by two virulence effectors, CodY and *rli60* [49,50]. Overall, our observations suggest that the adaptive response to BAC stress may influence virulence determinants. Alternatively, mutations introduced during mutagenesis that are not involved in BAC tolerance could account for the observed increase in hemolysis.

Taken together, our work suggests that the emergence of adaptive mutations for BAC tolerance in *L. monocytogenes* isolates found in food processing facilities could also result in strains that are more virulent (Figure 5) and have increased tolerance to particular antibiotics (Table 1, Figure 4). While the mutants we isolated did not develop BAC resistance at levels that would enable survival at BAC concentrations used for sanitization in industry, environments such as floor drains may present scenarios in which sanitizers are diluted to sub-lethal concentrations, resulting in continued exposure and subsequent adaptation. For instance, a recent study in *L. monocytogenes* observed that sub-lethal short-term exposure (10 days) to BAC and didecyldimethylammonium chloride resulted in decreased susceptibility to ciprofloxacin [25]. Additionally, several adaptive laboratory evolution studies in which *Pseudomonas aeruginosa* or *Escherichia coli* were grown in the presence of BAC have observed decreased sensitivity to BAC, and increased resistance to several unrelated antibiotics [74–76]. Thus, understanding the genetic underpinnings and consequences of BAC tolerance is vitally important for both controlling *L. monocytogenes* in the food processing environment and for preventing the spread of antibiotic resistance.

## Conflict of Interest

The authors declare that the research was conducted in the absence of any commercial or financial relationships that could be construed as a potential conflict of interest.

## Data Availability

*L. monocytogenes* FSL-N1-304 whole-genome Illumina and Oxford Nanopore Technologies sequencing data are available through the NCBI SRA run accession numbers SRR17006605 and SRR17067427, respectively. The FSL-N1-304 reference genome assembly is available through GenBank accession number GCA_021391455.1. Illumina whole-genome sequencing data for *mut-1* and *mut-2* are available through NCBI SRA run accession numbers SRR21641536 and SRR21641535, respectively.

## Funding

This material is based upon work supported by the National Institute of Food and Agriculture, U.S. Department of Agriculture, the Center for Agriculture, Food and the Environment, and the Food Science department at University of Massachusetts Amherst, under project number MAS-00529. The contents are solely the responsibility of the authors and do not necessarily represent the official views of the USDA or NIFA.

## Notes

### Competing Interest Statement

The authors have declared no competing interest.

## References

1. Ramaswamy V, Cresence VM, Rejitha JS, Lekshmi U, Dharsana KS, Prasad P, et al. Listeria-review of epidemiology and pathogenesis. J Microbiol Immunol Infect. 2007;40: 4–13.

2. Clauss HE, Lorber B. Central Nervous System Infection with Listeria monocytogenes. 2008 [cited 19 Jan 2023]. Available: http://www.cdc.gov/pulsenet/

3. Madjunkov M, Chaudhry S, Ito S. Listeriosis during pregnancy. Arch Gynecol Obstet. 296. doi:10.1007/s00404-017-4401-1

4. Scallan E, Hoekstra RM, Angulo FJ, Tauxe R V., Widdowson MA, Roy SL, et al. Foodborne Illness Acquired in the United States—Major Pathogens. Emerg Infect Dis. 2011;17: 7. doi:10.3201/EID1701.P11101

5. Townsend A, Strawn LK, Chapman BJ, Dunn LL. A Systematic Review of Listeria Species and Listeria monocytogenes Prevalence, Persistence, and Diversity throughout the Fresh Produce Supply Chain. Foods 2021, Vol 10, Page 1427. 2021;10: 1427. doi:10.3390/FOODS10061427

6. Farber JM, Zwietering M, Wiedmann M, Schaffner D, Hedberg CW, Harrison MA, et al. Alternative approaches to the risk management of Listeria monocytogenes in low risk foods. Food Control. 2021;123: 107601. doi:10.1016/J.FOODCONT.2020.107601

7. Pereira BMP, Tagkopoulos I. Benzalkonium Chlorides: Uses, Regulatory Status, and Microbial Resistance. Appl Environ Microbiol. 2019;85. doi:10.1128/AEM.00377-19

8. Wessels S, Ingmer H. Modes of action of three disinfectant active substances: a review. Regul Toxicol Pharmacol. 2013;67: 456–467. doi:10.1016/J.YRTPH.2013.09.006

9. Basaran P. Inhibition effect of belzalkonium chloride treatment on growth of common food contaminating fungal species. J Food Sci Technol. 2011;48: 515. doi:10.1007/S13197-011-0268-5

10. Gelbicova T, Florianova M, Hluchanova L, Kalova A, Korena K, Strakova N, et al. Comparative Analysis of Genetic Determinants Encoding Cadmium, Arsenic, and Benzalkonium Chloride Resistance in Listeria monocytogenes of Human, Food, and Environmental Origin. Front Microbiol. 2021;11: 3397. doi:10.3389/FMICB.2020.599882/BIBTEX

11. Meier AB, Guldimann C, Markkula A, Pöntinen A, Korkeala H, Tasara T. Comparative Phenotypic and Genotypic Analysis of Swiss and Finnish Listeria monocytogenes Isolates with Respect to Benzalkonium Chloride Resistance. Front Microbiol. 2017;8. doi:10.3389/FMICB.2017.00397

12. Ratani SS, Siletzky RM, Dutta V, Yildirim S, Osborne JA, Lin W, et al. Heavy metal and disinfectant resistance of Listeria monocytogenes from foods and food processing plants. Appl Environ Microbiol. 2012;78: 6938–6945. doi:10.1128/AEM.01553-12/ASSET/3774F514-37AE-4510-8550-2A8828C53EAA/ASSETS/GRAPHIC/ZAM9991037010004.JPEG

13. Romanova NA, Wolffs PFG, Brovko LY, Griffiths MW. Role of efflux pumps in adaptation and resistance of Listeria monocytogenes to benzalkonium chloride. Appl Environ Microbiol. 2006;72: 3498–3503. doi:10.1128/AEM.72.5.3498-3503.2006/ASSET/FBB76CE4-0E7C-4440-9640-CA891E358D69/ASSETS/GRAPHIC/ZAM0050667720001.JPEG

14. Soumet C, Ragimbeau C, Maris P. Screening of benzalkonium chloride resistance in Listeria monocytogenes strains isolated during cold smoked fish production. Lett Appl Microbiol. 2005;41: 291–296. doi:10.1111/J.1472-765X.2005.01763.X

15. Xu D, Li Y, Shamim Hasan Zahid M, Yamasaki S, Shi L, Li Jrong, et al. Benzalkonium chloride and heavy-metal tolerance in Listeria monocytogenes from retail foods. Int J Food Microbiol. 2014;190: 24–30. doi:10.1016/J.IJFOODMICRO.2014.08.017

16. Hegstad K, Langsrud S, Lunestad BT, Scheie AA, Sunde M, Yazdankhah SP. Does the wide use of quaternary ammonium compounds enhance the selection and spread of antimicrobial resistance and thus threaten our health? Microb Drug Resist. 2010;16: 91–104. doi:10.1089/MDR.2009.0120

17. Mc Carlie S, Boucher CE, Bragg RR. Molecular basis of bacterial disinfectant resistance. Drug Resist Updat. 2020;48. doi:10.1016/J.DRUP.2019.100672

18. To MS, Favrin S, Romanova N, Griffiths MW. Postadaptational resistance to benzalkonium chloride and subsequent physicochemical modifications of Listeria monocytogenes. Appl Environ Microbiol. 2002;68: 5258–5264. doi:10.1128/AEM.68.11.5258-5264.2002/ASSET/BD247714-A39D-456A-ABDE-43B7820D7A66/ASSETS/GRAPHIC/AM1120249001.JPEG

19. Yu T, Jiang X, Zhang Y, Ji S, Gao W, Shi L. Effect of Benzalkonium Chloride adaptation on sensitivity to antimicrobial agents and tolerance to environmental stresses in Listeria monocytogenes. Front Microbiol. 2018;9: 2906. doi:10.3389/FMICB.2018.02906/BIBTEX

20. Jiang X, Yu T, Xu Y, Wang H, Korkeala H, Shi L. MdrL, a major facilitator superfamily efflux pump of Listeria monocytogenes involved in tolerance to benzalkonium chloride. Appl Microbiol Biotechnol. 2019;103: 1339–1350. doi:10.1007/S00253-018-9551-Y/FIGURES/6

21. Guérin F, Galimand M, Tuambilangana F, Courvalin P, Cattoir V. Overexpression of the Novel MATE Fluoroquinolone Efflux Pump FepA in Listeria monocytogenes Is Driven by Inactivation of Its Local Repressor FepR. PLoS One. 2014;9: e106340. doi:10.1371/JOURNAL.PONE.0106340

22. Douarre PE, Sévellec Y, Le Grandois P, Soumet C, Bridier A, Roussel S. FepR as a Central Genetic Target in the Adaptation to Quaternary Ammonium Compounds and Cross-Resistance to Ciprofloxacin in Listeria monocytogenes. Front Microbiol. 2022;13: 864576. doi:10.3389/FMICB.2022.864576/BIBTEX

23. Kode D, Nannapaneni R, Bansal M, Chang S, Cheng WH, Sharma CS, et al. Low-level tolerance to fluoroquinolone antibiotic ciprofloxacin in qac-adapted subpopulations of listeria monocytogenes. Microorganisms. 2021;9: 1052. doi:10.3390/MICROORGANISMS9051052/S1

24. Buffet-Bataillon S, Tattevin P, Maillard JY, Bonnaure-Mallet M, Jolivet-Gougeon A. Efflux pump induction by quaternary ammonium compounds and fluoroquinolone resistance in bacteria. Future Microbiol. 2016;11: 81–92. doi:10.2217/FMB.15.131

25. Guérin A, Bridier A, Le Grandois P, Sévellec Y, Palma F, Félix B, et al. Exposure to Quaternary Ammonium Compounds Selects Resistance to Ciprofloxacin in Listeria monocytogenes. Pathogens 2021, Vol 10, Page 220. 2021;10: 220. doi:10.3390/PATHOGENS10020220

26. Norton DM, Scarlett JM, Horton K, Sue D, Thimothe J, Boor KJ, et al. Characterization and Pathogenic Potential of Listeria monocytogenes Isolates from the Smoked Fish Industry. Appl Environ Microbiol. 2001;67: 646. doi:10.1128/AEM.67.2.646-653.2001

27. Bose JL. Chemical and UV Mutagenesis. Methods Mol Biol. 2016;1373: 111–115. doi:10.1007/7651_2014_190

28. Haney EF, Trimble MJ, Hancock REW. Microtiter plate assays to assess antibiofilm activity against bacteria. Nature Protocols 2021 16:5. 2021;16: 2615–2632. doi:10.1038/s41596-021-00515-3

29. SAMPATHKUMAR B, TSOUGRIANI E, YU LSL, KHACHATOURIANS GG. A quantitative microtiter plate hemolysis assay for Listeria monocytogenes. J Food Saf. 1998;18: 197–203. doi:10.1111/j.1745-4565.1998.tb00214.x

30. Clinical and Laboratory Standards Institute. CLSI. Performance Standards for Antimicrobial Susceptibility Testing. 32nd ed. CLSI supplement M100. 32nd ed. 2022.

31. bcl2fastq: A proprietary Illumina software for the conversion of bcl files to basecalls. https://support.illumina.com/sequencing/sequencing_software/bcl2fastq-conversion-software.html;

32. Wick RR, Judd LM, Gorrie CL, Holt KE. Unicycler: Resolving bacterial genome assemblies from short and long sequencing reads. PLoS Comput Biol. 2017;13: e1005595. doi:10.1371/journal.pcbi.1005595

33. Gurevich A, Saveliev V, Vyahhi N, Tesler G. QUAST: quality assessment tool for genome assemblies. Bioinformatics. 2013;29: 1072–1075. doi:10.1093/bioinformatics/btt086

34. Seemann T. Prokka: rapid prokaryotic genome annotation. Bioinformatics. 2014;30: 2068–2069. doi:10.1093/BIOINFORMATICS/BTU153

35. Martin M. Cutadapt removes adapter sequences from high-throughput sequencing reads. EMBnet J. 2011;17: 10–12. doi:10.14806/EJ.17.1.200

36. Krueger F. “Trim galore” A wrapper tool around Cutadapt and FastQC to consistently apply quality and adapter trimming to FastQ files. 2015; 516.

37. Li H. Aligning sequence reads, clone sequences and assembly contigs with BWA-MEM. 2013 [cited 29 Jun 2022]. doi:10.48550/arxiv.1303.3997

38. Danecek P, Bonfield JK, Liddle J, Marshall J, Ohan V, Pollard MO, et al. Twelve years of SAMtools and BCFtools. Gigascience. 2021;10. doi:10.1093/GIGASCIENCE/GIAB008

39. Garrison E, Marth G. Haplotype-based variant detection from short-read sequencing. 2012.

40. McKenna A, Hanna M, Banks E, Sivachenko A, Cibulskis K, Kernytsky A, et al. The Genome Analysis Toolkit: a MapReduce framework for analyzing next-generation DNA sequencing data. Genome Res. 2010;20: 1297–1303. doi:10.1101/GR.107524.110

41. Cingolani P, Platts A, Wang LL, Coon M, Nguyen T, Wang L, et al. A program for annotating and predicting the effects of single nucleotide polymorphisms, SnpEff: SNPs in the genome of Drosophila melanogaster strain w1118; iso-2; iso-3. Fly (Austin). 2012;6: 80–92. doi:10.4161/FLY.19695

42. Sahl JW, Gregory Caporaso J, Rasko DA, Keim P. The large-scale blast score ratio (LS-BSR) pipeline: a method to rapidly compare genetic content between bacterial genomes. PeerJ. 2014;2. doi:10.7717/PEERJ.332

43. Mitchell AL, Attwood TK, Babbitt PC, Blum M, Bork P, Bridge A, et al. InterPro in 2019: improving coverage, classification and access to protein sequence annotations. Nucleic Acids Res. 2019;47: D351–D360. doi:10.1093/NAR/GKY1100

44. Quinlan AR, Hall IM. BEDTools: a flexible suite of utilities for comparing genomic features. Bioinformatics. 2010;26: 841–842. doi:10.1093/BIOINFORMATICS/BTQ033

45. Love MI, Huber W, Anders S. Moderated estimation of fold change and dispersion for RNA-seq data with DESeq2. Genome Biol. 2014;15: 1–21. doi:10.1186/S13059-014-0550-8/FIGURES/9

46. SAS Institute Inc. JMP® Pro. Cary, NC; pp. 1989–2023.

47. Wickham H. ggplot2: Elegant Graphics for Data Analysis. Springer-Verlag. 2016.

48. Kanehisa M, Goto S. KEGG: kyoto encyclopedia of genes and genomes. Nucleic Acids Res. 2000;28: 27–30. doi:10.1093/NAR/28.1.27

49. Marinho CM, Dos Santos PT, Kallipolitis BH, Johansson J, Ignatov D, Guerreiro DN, et al. The σB-dependent regulatory sRNA Rli47 represses isoleucine biosynthesis in Listeria monocytogenes through a direct interaction with the ilvA transcript. RNA Biol. 2019;16: 1424–1437. doi:10.1080/15476286.2019.1632776

50. Brenner M, Lobel L, Borovok I, Sigal N, Herskovits AA. Controlled branched-chain amino acids auxotrophy in Listeria monocytogenes allows isoleucine to serve as a host signal and virulence effector. PLoS Genet. 2018;14: e1007283. doi:10.1371/JOURNAL.PGEN.1007283

51. Dutta V, Elhanaf D, Kathariou S. Conservation and Distribution of the Benzalkonium Chloride Resistance Cassette bcrABC in Listeria monocytogenes. Appl Environ Microbiol. 2013;79: 6067–6074. doi:10.1128/AEM.01751-13/ASSET/67B5E713-5DB2-4AE5-901F-C4A365161EBE/ASSETS/GRAPHIC/ZAM9991047400003.JPEG

52. Elhanafi D, Utta V, Kathariou S. Genetic characterization of plasmid-associated benzalkonium chloride resistance determinants in a listeria monocytogenes train from the 1998 -1999 outbreak. Appl Environ Microbiol. 2010;76: 8231–8238. doi:10.1128/AEM.02056-10/ASSET/4E7916C1-4D47-44D6-997A-D8EC8CB7E6FC/ASSETS/GRAPHIC/ZAM9991016160005.JPEG

53. Müller A, Rychli K, Muhterem-Uyar M, Zaiser A, Stessl B, Guinane CM, et al. Tn6188 -A Novel Transposon in Listeria monocytogenes Responsible for Tolerance to Benzalkonium Chloride. PLoS One. 2013;8: e76835. doi:10.1371/JOURNAL.PONE.0076835

54. Chen GY, Kao CY, Smith HB, Rust DP, Powers ZM, Li AY, et al. Mutation of the Transcriptional Regulator YtoI Rescues Listeria monocytogenes Mutants Deficient in the Essential Shared Metabolite 1,4-Dihydroxy-2-Naphthoate (DHNA). Infect Immun. 2020;88. doi:10.1128/IAI.00366-19

55. Kelliher JL, Grunenwald CM, Abrahams RR, Daanen ME, Lew CI, Rose WE, et al. PASTA kinase-dependent control of peptidoglycan synthesis via ReoM is required for cell wall stress responses, cytosolic survival, and virulence in Listeria monocytogenes. PLoS Pathog. 2021;17: e1009881. doi:10.1371/JOURNAL.PPAT.1009881

56. Walsh JP, Bell RM. Diacylglycerol kinase from Escherichia coli. 1992. pp. 153–162. doi:10.1016/0076-6879(92)09019-Y

57. Park YK, Lee JY, Ko KS. Transcriptomic analysis of colistin-susceptible and colistin-resistant isolates identifies genes associated with colistin resistance in Acinetobacter baumannii. Clinical Microbiology and Infection. 2015;21: 765.e1–765.e7. doi:10.1016/J.CMI.2015.04.009

58. Guinane CM, Cotter PD, Ross RP, Hill C. Contribution of penicillin-binding protein homologs to antibiotic resistance, cell morphology, and virulence of Listeria monocytogenes EGDe. Antimicrob Agents Chemother. 2006;50: 2824–2828. doi:10.1128/AAC.00167-06/ASSET/D73488E3-F530-4EF5-8691-18A324495231/ASSETS/GRAPHIC/ZAC0080659680002.JPEG

59. McKay SL, Portnoy DA. Ribosome hibernation facilitates tolerance of stationary-phase bacteria to aminoglycosides. Antimicrob Agents Chemother. 2015;59: 6992–6999. doi:10.1128/AAC.01532-15/ASSET/152CF37F-F3F3-4054-8A69-6A35FCC0258A/ASSETS/GRAPHIC/ZAC0111545370004.JPEG

60. Kremer PHC, Lees JA, Koopmans MM, Ferwerda B, Arends AWM, Feller MM, et al. Benzalkonium tolerance genes and outcome in Listeria monocytogenes meningitis. Clinical Microbiology and Infection. 2017;23: 265.e1–265.e7. doi:10.1016/J.CMI.2016.12.008

61. Bolten S, Harrand AS, Skeens J, Wiedmann M. Nonsynonymous Mutations in fepR Are Associated with Adaptation of Listeria monocytogenes and Other Listeria spp. to Low Concentrations of Benzalkonium Chloride but Do Not Increase Survival of L. monocytogenes and Other Listeria spp. after Exposure to Benzalkonium Chloride Concentrations Recommended for Use in Food Processing Environments. Appl Environ Microbiol. 2022;88. doi:10.1128/AEM.00486-22/ASSET/C7691245-6758-41F6-8274-58D7CF1C90A0/ASSETS/IMAGES/LARGE/AEM.00486-22-F003.JPG

62. Hussein SH, Samir R, Aziz RK, Toama MA. Two putative MmpL homologs contribute to antimicrobial resistance and nephropathy of enterohemorrhagic E. coli O157:H7. Gut Pathog. 2019;11: 1–13. doi:10.1186/S13099-019-0296-7/TABLES/1

63. Bowman JP, Hages E, Nilsson RE, Kocharunchitt C, Ross T. Investigation of the listeria monocytogenes Scott A acid tolerance response and associated physiological and phenotypic features via whole proteome analysis. J Proteome Res. 2012;11: 2409–2426. doi:10.1021/PR201137C/SUPPL_FILE/PR201137C_SI_008.PDF

64. Reith J, Mayer C. Peptidoglycan turnover and recycling in Gram-Positive bacteria. Appl Microbiol Biotechnol. 2011;92: 1–11. doi:10.1007/S00253-011-3486-X/FIGURES/2

65. Fox EM, Leonard N, Jordan K. Physiological and transcriptional characterization of persistent and nonpersistent Listeria monocytogenes isolates. Appl Environ Microbiol. 2011;77: 6559–6569. doi:10.1128/AEM.05529-11/ASSET/E53F4AC5-6AF2-4E12-A8F2-C08C3D55F894/ASSETS/GRAPHIC/ZAM9991024670003.JPEG

66. Casey A, Fox EM, Schmitz-Esser S, Coffey A, Mcauliffe O, Jordan K, et al. Transcriptome analysis of Listeria monocytogenes exposed to biocide stress reveals a multi-system response involving cell wall synthesis, sugar uptake, and motility. 2014. doi:10.3389/fmicb.2014.00068

67. Dalet K, Cenatiempo Y, Cossart P, Glaser P, Amend A, Baquero-Mochales F, et al. A σ54-dependent PTS permease of the mannose family is responsible for sensitivity of Listeria monocytogenes to mesentericin Y105. Microbiology (N Y). 2001;147: 3263–3269. doi:10.1099/00221287-147-12-3263/CITE/REFWORKS

68. Xue J, Hunter I, Steinmetz T, Peters A, Ray B, Miller KW. Novel activator of mannose-specific phosphotransferase system permease expression in Listeria innocua, identified by screening for pediocin AcH resistance. Appl Environ Microbiol. 2005;71: 1283–1290. doi:10.1128/AEM.71.3.1283-1290.2005/ASSET/42BA2BFF-CD9E-4E2D-BA4E-3A877477275E/ASSETS/GRAPHIC/ZAM0030552640004.JPEG

69. Geldart K, Kaznessis YN. Characterization of class IIa bacteriocin resistance in Enterococcus faecium. Antimicrob Agents Chemother. 2017;61. doi:10.1128/AAC.02033-16/SUPPL_FILE/ZAC004176031S1.PDF

70. Vadyvaloo V, Arous S, Gravesen A, Héchard Y, Chauhan-Haubrock R, Hastings JW, et al. Cell-surface alterations in class IIa bacteriocin-resistant Listeria monocytogenes strains. Microbiology (N Y). 2004;150: 3025–3033. doi:10.1099/MIC.0.27059-0/CITE/REFWORKS

71. Personnic N, Bruck S, Nahori MA, Toledo-Arana A, Nikitas G, Lecuit M, et al. The Stress-Induced Virulence Protein InlH Controls Interleukin-6 Production during Murine Listeriosis. Infect Immun. 2010;78: 1979. doi:10.1128/IAI.01096-09

72. Rouquette C, Ripio MT, Pellegrini E, Bolla JM, Tascon RI, Vázquez-Boland JA, et al. Identification of a ClpC ATPase required for stress tolerance and in vivo survival of Listeria monocytogenes. Mol Microbiol. 1996;21: 977–987. doi:10.1046/J.1365-2958.1996.641432.X

73. Rouquette C, De Chastellier C, Nair S, Berche P. The ClpC ATPase of Listeria monocytogenes is a general stress protein required for virulence and promoting early bacterial escape from the phagosome of macrophages. Mol Microbiol. 1998;27: 1235– 1245. doi:10.1046/J.1365-2958.1998.00775.X

74. Mc Cay PH, Ocampo-Sosa AA, Fleming GTA. Effect of subinhibitory concentrations of benzalkonium chloride on the competitiveness of Pseudomonas aeruginosa grown in continuous culture. Microbiology (Reading). 2010;156: 30–38. doi:10.1099/MIC.0.029751-0

75. Nordholt N, Kanaris O, Schmidt SBI, Schreiber F. Persistence against benzalkonium chloride promotes rapid evolution of tolerance during periodic disinfection. Nat Commun. 2021;12. doi:10.1038/S41467-021-27019-8

76. Tandukar M, Oh S, Tezel U, Konstantinidis KT, Pavlostathis SG. Long-term exposure to benzalkonium chloride disinfectants results in change of microbial community structure and increased antimicrobial resistance. Environ Sci Technol. 2013;47: 9730–9738. doi:10.1021/ES401507K

