## Supplementary figures and images for "Chemical mutagenesis of *Listeria monocytogenes* for increased tolerance to benzalkonium chloride shows independent genetic underpinnings and off-target antibiotic resistance"

### Supplemental Table S1

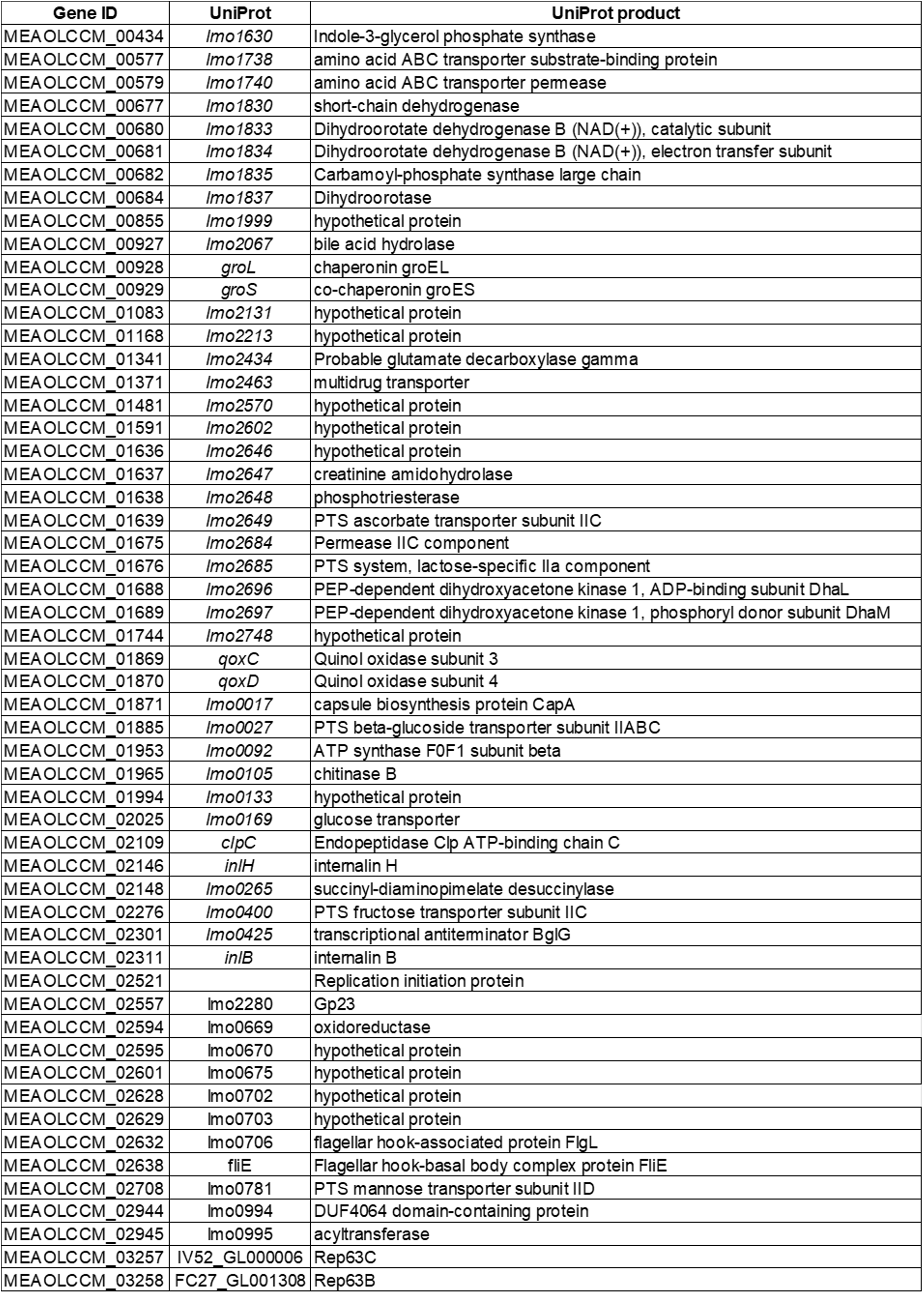

### Supplemental Table S2

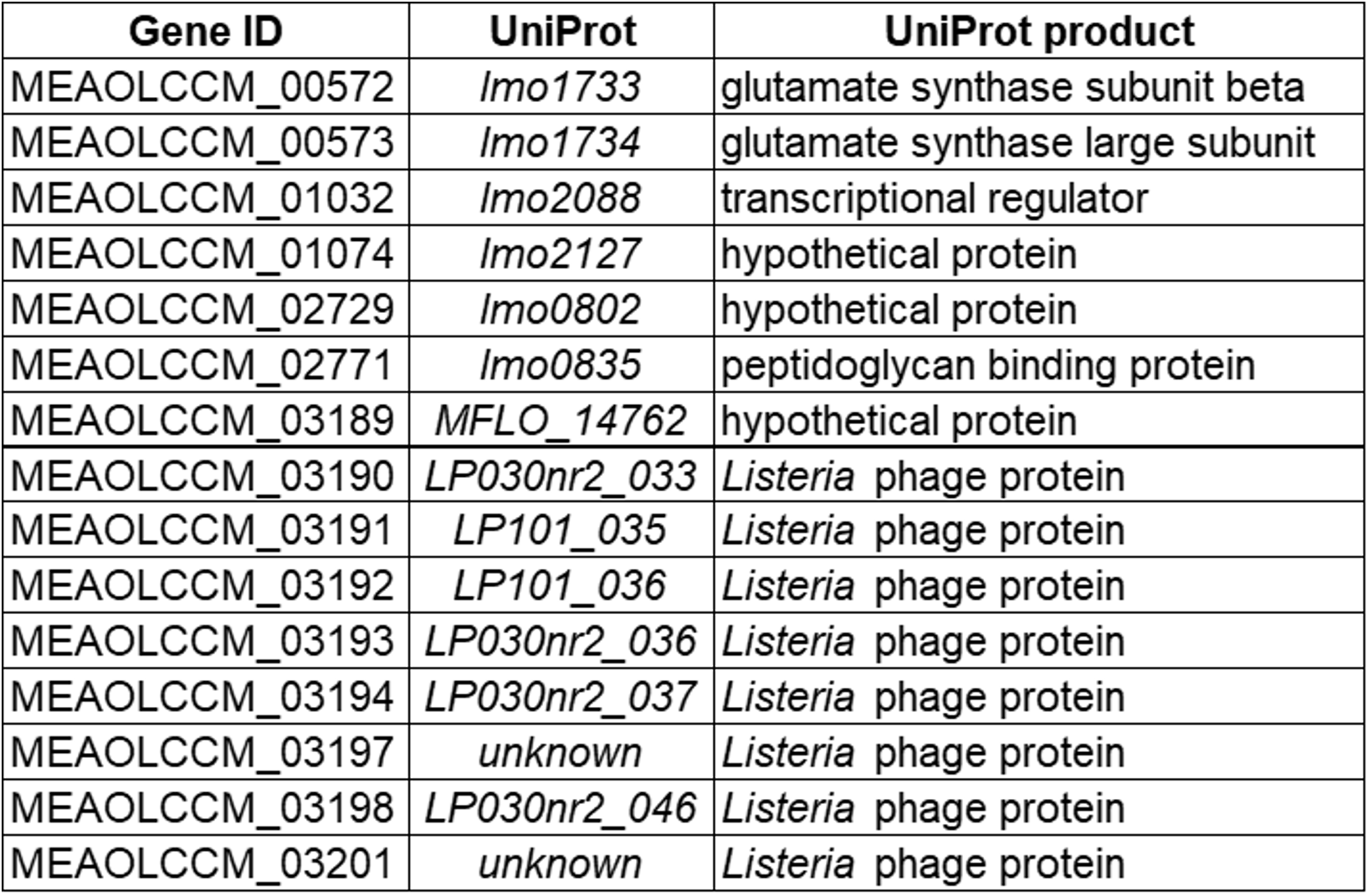

### Supplemental Table S3

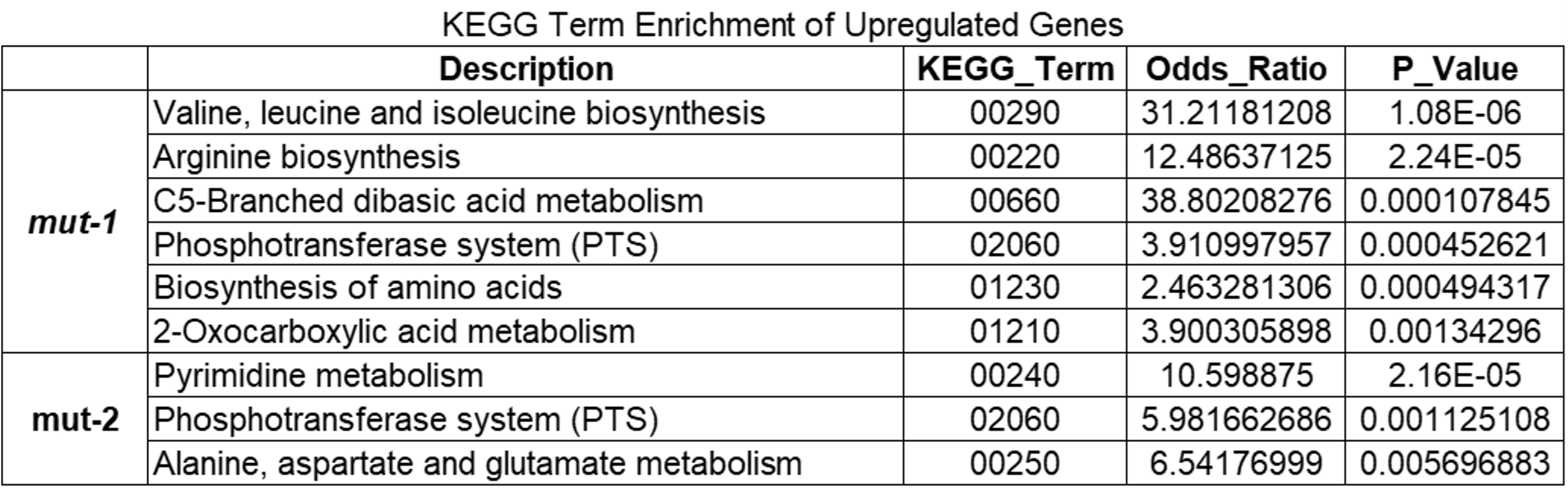

### Supplemental Table S4

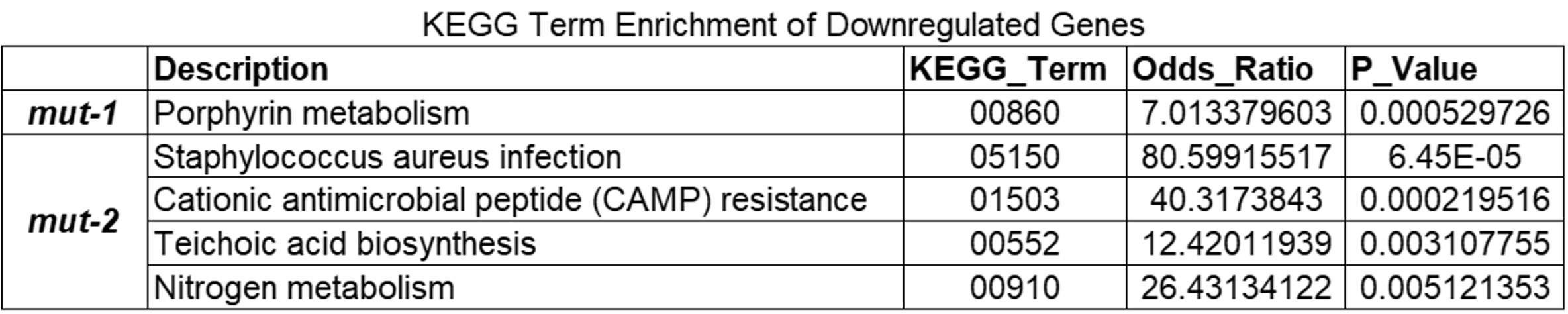
